# Nuclear Pore Transport: Toward an Integrated Perspective

**DOI:** 10.1101/2025.09.18.677018

**Authors:** Luca Lanzano, Francesco Cardarelli

## Abstract

Nuclear pore complexes (NPCs) mediate the selective exchange of proteins and RNAs between nucleus and cytoplasm. Despite decades of study, the molecular mechanism of transport remains debated, particularly the role of FG-nucleoporin dynamics in facilitating translocation. Here, we compare high-resolution single-molecule trajectories obtained with MINFLUX microscopy with earlier correlation-based measurements, revealing that actual pore-crossing events are typically completed within 2-4 localization steps, corresponding to 2-6 ms depending on localization rates. These values closely match transit times obtained from fluorescence fluctuation analysis and align with models in which cargo binding induces transient FG-nucleoporin collapse. Together with recent independent observations, this synthesis supports a converging view of NPC transport as a fast, directional, FG-assisted process. We propose that integrating fluctuation-based and trajectory-based approaches provides a robust framework for reconciling mechanistic hypotheses and refining our understanding of selective gating.

## Introduction: the unresolved question of nuclear transport dynamics

Nucleocytoplasmic transport is essential for eukaryotic life, mediating the bidirectional exchange of proteins and RNA through nuclear pore complexes (NPCs)—massive assemblies embedded in the nuclear envelope that maintain the selectivity barrier between nucleus and cytoplasm^1,2^. Despite decades of research, the mechanism enabling fast and selective transport of cargoes remains largely unresolved. The debate centers on a single overarching question concerning the physical nature of the permeability barrier—do translocating molecules cross the NPC purely by diffusion, or rather via transient, directional interactions with FG-nucleoporins (FG-Nups)?

These two translocation mechanisms correspond to two major experimental models that propose distinct structural organizations of FG-Nups within the NPC. The first envisions permeation as a diffusive process: FG-repeat domains assemble into a cohesive, gel-like mesh that acts as an entropic sieve, allowing receptor-bound cargos to pass while excluding inert macromolecules^3^. This idea was refined in the selective phase model, which proposed that FG domains organize into a polymeric network forming a thermodynamic barrier against unspecific passage, while still enabling passive, diffusion-driven translocation of cargos^4,5^. By contrast, a second model posits a directed mode of transport. In this framework, the FG-Nups within the permeability barrier undergo local rearrangements upon receptor binding and transiently collapse to create directional conduits that guide cargos across the pore^6,7^. Experimental support for this mechanism comes from in vitro observations—including AFM and EM evidence of receptor-induced FG condensation^6,7^—as well as from live-cell fluorescence correlation spectroscopy (FCS), which detected rapid and directional transport signatures at the single-pore level^8,9^.

Over the past two decades, single-particle tracking (SPT) has provided foundational insights into nucleocytoplasmic transport. Early live-cell SPT studies, including those by Grünwald, Musser, and colleagues, demonstrated that individual cargos could be followed as they approached and entered the NPC, providing dwell time distributions and revealing heterogeneous transport behavior^10-15^. These experiments firmly established that transport could be studied at the single-molecule level in intact cells, even if spatial precision and temporal resolution were not yet sufficient to unambiguously resolve the millisecond dynamics of barrier crossing. The challenge, however, was not methodological but intrinsic: capturing the fast, confined motion within the ~50-nm central channel.

Recent breakthroughs in single-molecule imaging now allow the field to revisit this question. Most notably, MINFLUX nanoscopy now enables the tracking of individual cargos traversing NPCs in intact cells with nanometer spatial precision and millisecond temporal resolution^16^, offering an unprecedented opportunity to directly confront existing models with observation.

### The renewed debate: insights from MINFLUX and complementary studies

In a recent *Nature* study, Sau et al. applied 3D MINFLUX microscopy to visualize nuclear transport in living cells, achieving unmatched spatial and temporal accuracy^16^. Their trajectories reveal that import and export occur along overlapping annular pathways within the NPC and that cargos remain within a region defined as −25 to +25 nm from the midplane for an average of ~14 ms. These findings represent a technical milestone, adding valuable geometric detail to the long-standing question of transport routes.

However, whether this “residence time” corresponds to the actual translocation step is uncertain. Examination of the authors’ published trajectories and videos suggests that the true crossing—from one side of the pore to the other across the central plane—is completed in far fewer steps than implied by the full ±25 nm range. Rather than a prolonged random walk, the transition appears as a short, directional burst embedded within a broader exploratory phase.

Independent evidence for directed motion has recently emerged from cargo-centric tracking approaches. Li et al., in a 2024 *Nature Physics* article, reported that nuclear transport receptors exhibit anisotropic, non-random trajectories during NPC crossing^17^. Their results strongly support the concept of biased progression along defined paths rather than unrestricted Brownian diffusion. Taken together, these studies underscore that the NPC does not behave as a passive channel but as a dynamic structure that constrains and facilitates passage once productive interactions are established.

### Revisiting early evidence: fluctuation analysis of NPC transport

More than a decade ago, we introduced a method that combined orbital tracking of individual NPCs with fluorescence fluctuation and cross-correlation analysis to probe molecular dynamics without reconstructing trajectories^8,9^. This approach exploited intensity fluctuations along a nanometric orbit encircling a single pore and identified directed transport statistically, using pair-correlation of signals recorded on opposite sides of the channel.

This analysis yielded two key insights. First, transit times derived from multiple events showed sharply peaked distributions centered at 3–5 ms, implying a narrowly defined kinetic step for barrier crossing. Second, cross-correlation between cargo and Nup153 revealed synchronous fluctuations, suggesting transient co-movement, compatible with receptor-induced rearrangements of FG domains. These findings supported the mechanistic model proposed by Lim et al.^6,7^, in which binding of transport receptors induces FG domain collapse, generating a directional bias that facilitates selective transport. Although complementary to earlier SPT studies, our approach was distinctive in detecting transport signatures statistically—at 1-ms temporal resolution—without trajectory reconstruction. This latter difference also explains why some skepticism remained: correlation analysis, while powerful, does not provide direct visualization of individual trajectories. The advent of MINFLUX offers the opportunity to validate, at the single-trajectory level, the mechanistic insights previously inferred from fluctuation analysis.

### Re-analysis of MINFLUX trajectories: fast and directional crossing

To test whether MINFLUX data align with correlation-based predictions, we re-examined representative import and export trajectories from Sau et al.’s supplementary movies. For each, we isolated the segment corresponding to true translocation—defined as the crossing of the 0-nm plane from one side of the NPC to the other—and mapped these localizations onto the observation geometry employed in our earlier orbital tracking experiments (Figures 1 and 2). This alignment enables direct comparison between correlation-based inference and high-resolution imaging.

**Figure 1.**
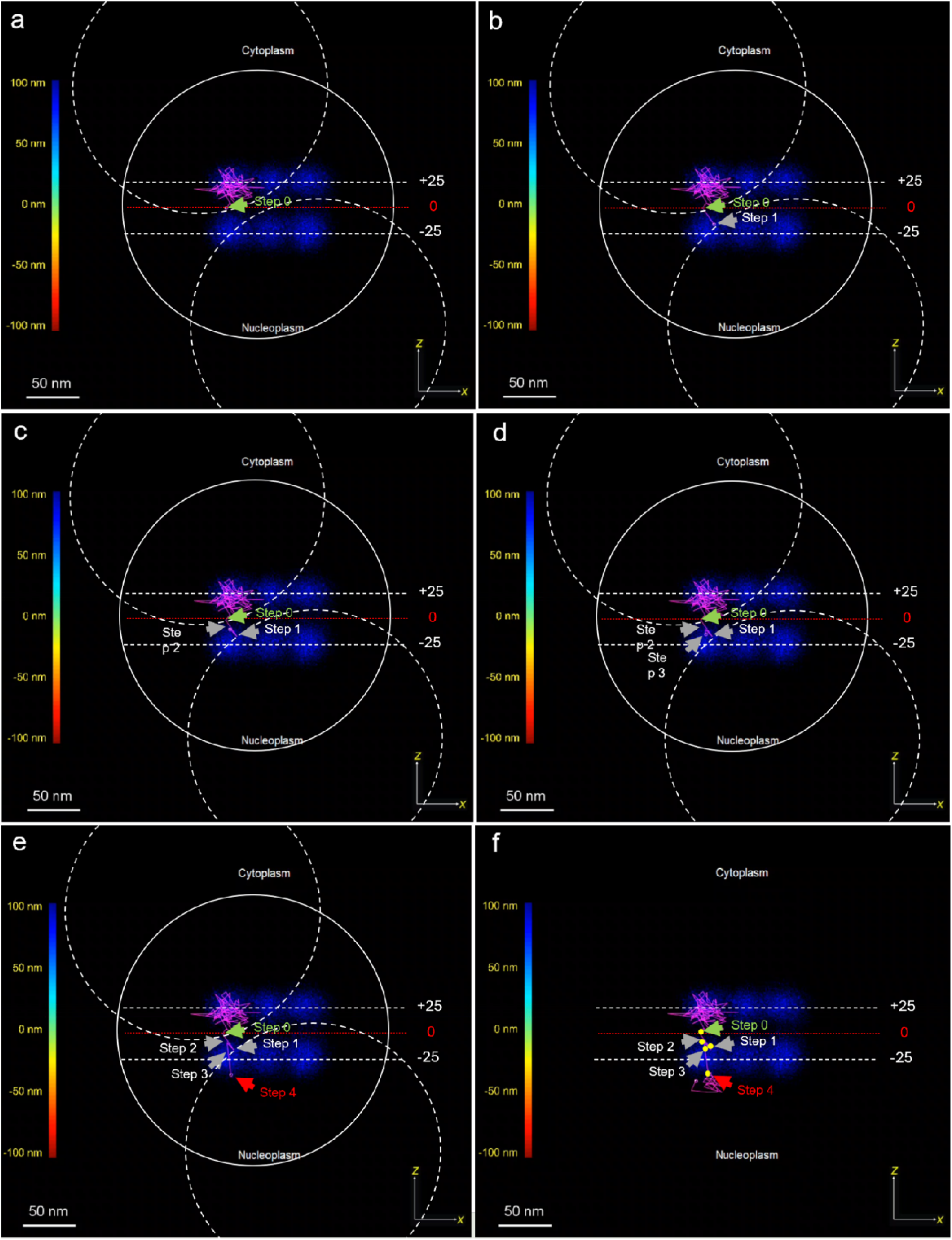
Single import trajectory overlaid with orbital tracking geometry. Representative import trajectory extracted from Supplementary Movie 1 of Sau et al. (2025), displayed across selected frames (a–f). The nanometric observation geometry used in Cardarelli et al. (2012) is superimposed to illustrate the region probed by pair-correlation. Gray discs indicate point spread functions corresponding to correlation sites. White dashed lines mark ±25 nm boundaries from the pore center (0 nm, red). Four consecutive localizations (green to red arrows) span the crossing event. Assuming localization intervals of 0.52–1.6 ms, this corresponds to a transit time of ~2–6 ms, matching correlation-based estimates.

**Figure 2.**
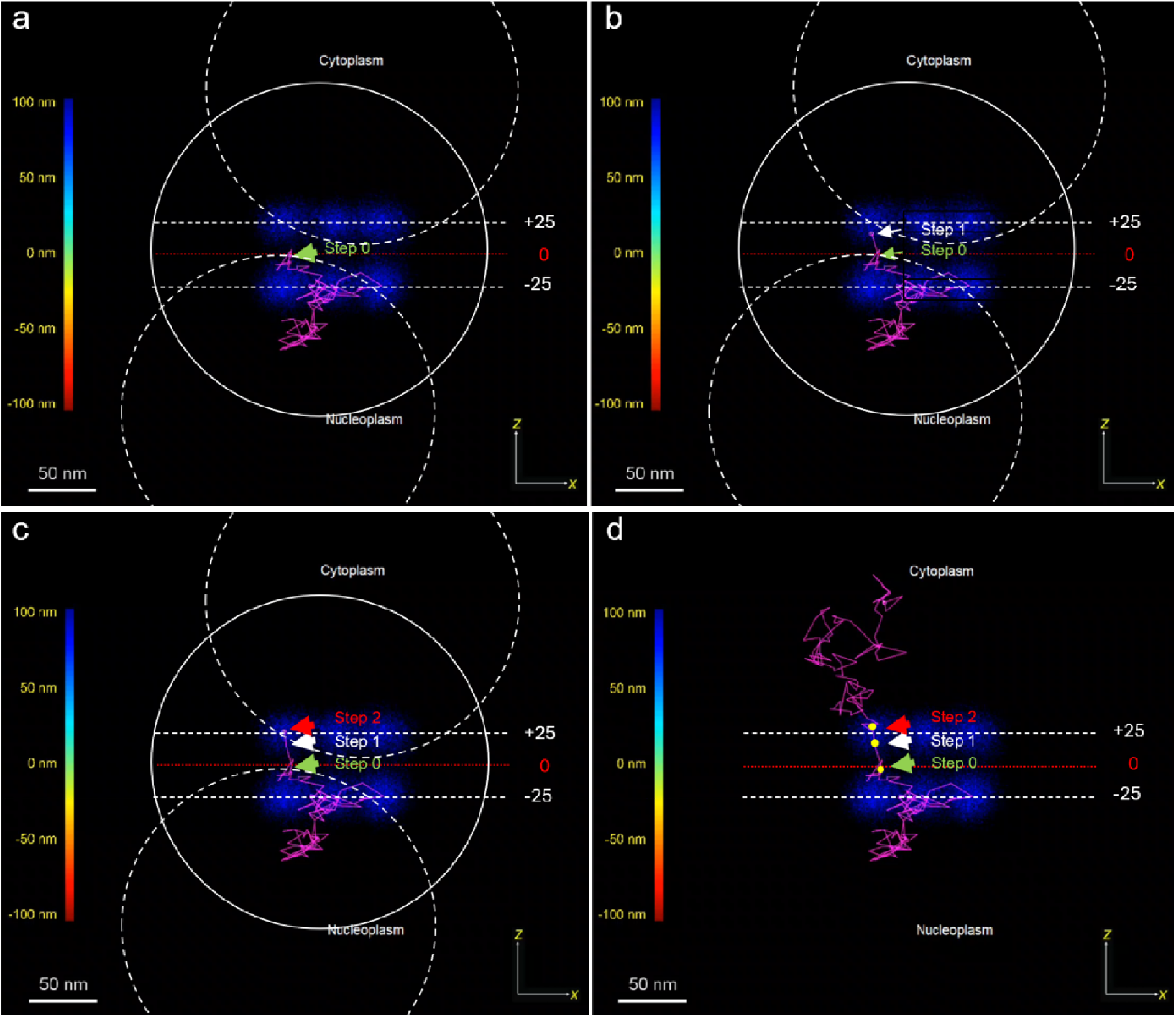
Single export trajectory overlaid with orbital tracking geometry. As in Figure 1, but for an export event extracted from Supplementary Movie 2 of Sau et al. (2025). The translocation segment comprises only two consecutive steps, yielding an estimated crossing time of ~1–3 ms. These values reinforce the convergence between MINFLUX tracking and fluctuation-derived kinetics.

The analysis highlights two consistent features. First, the active crossing phase typically involves only 2–4 consecutive localizations, far fewer than suggested by the broader ±25 nm interval. Second, applying Sau et al.’s reported localization intervals (0.52 ms minimum, 1.6 ms average) yields estimated crossing durations of ~2–6 ms for import and ~1–3 ms for export— values that fall squarely within the distributions we reported over a decade ago^8,9^. Figures 1 and 2 illustrate these findings, showing how the mechanistic signal extracted from fluctuation analysis corresponds to a brief, directional event now directly observable with MINFLUX.

### Toward a unified mechanistic framework

The convergence of fluctuation spectroscopy, MINFLUX imaging, and cargo-centric analysis strengthens the case for a model in which FG-Nups act as dynamic, receptor-responsive polymers. This view is consistent with nanomechanical data showing receptor-induced FG condensation and with structural models proposing reversible collapse as a gating principle^6,7,11^. Directional bias may emerge from sequential stabilization of receptor– FG contacts, enabling “hand-over-hand” translocation that reconciles selectivity with high speed.

These considerations help redefine the priorities for experimental design. Future studies should integrate high-speed, multi-color tracking with probes reporting FG-Nup conformation, alongside statistical approaches capable of resolving microsecond fluctuations. Such integrative strategies will be essential to capture the interplay of molecular recognition, polymer dynamics, and confinement that underpins the NPC’s extraordinary performance.

More than a decade ago, correlation analysis suggested that NPC transport is fast, directional, and FG-mediated^8,9^. Today, MINFLUX and complementary approaches provide direct support for this interpretation^16,17^. Far from being contradictory, these methods converge on a coherent picture: the NPC is not a static filter but a responsive molecular machine. The challenge ahead lies in integrating these insights into a comprehensive, experimentally grounded model of selective gating.

## Acknowledgments

We are deeply grateful to Enrico Gratton, whose mentorship and vision shaped the foundations of this work.

